# Hormonal steroids bind the *Neisseria gonorrhoeae* multidrug resistance regulator, MtrR, to induce a multidrug binding efflux pump and stress-response sigma factor

**DOI:** 10.1101/2023.06.13.544409

**Authors:** Grace M. Hooks, Julio C. Ayala, Grace A. Beggs, John R. Perfect, Maria A. Schumacher, William M. Shafer, Richard G. Brennan

## Abstract

Overexpression of the multidrug efflux pump MtrCDE, a critical factor of multidrug-resistance in *Neisseria gonorrhoeae*, the causative agent of gonorrheae, is repressed by the transcriptional regulator, MtrR (multiple transferable resistance repressor). Here, we report the results from a series of *in vitro* experiments to identify innate, human inducers of MtrR and to understand the biochemical and structural mechanisms of the gene regulatory function of MtrR. Isothermal titration calorimetry experiments reveal that MtrR binds the hormonal steroids progesterone, β-estradiol, and testosterone, all of which are present at significant concentrations at urogenital infection sites as well as ethinyl estrogen, a component of some birth control pills. Binding of these steroids results in decreased affinity of MtrR for cognate DNA, as demonstrated by fluorescence polarization-based assays. The crystal structures of MtrR bound to each steroid provided insight into the flexibility of the binding pocket, elucidated specific residue-ligand interactions, and revealed the conformational consequences of the induction mechanism of MtrR. Three residues, D171, W136 and R176 are key to the specific binding of these gonadal steroids. These studies provide a molecular understanding of the transcriptional regulation by MtrR that promotes *N. gonorrhoeae* survival in its human host.

## Introduction

The pathogen *Neisseria gonorrhoeae* is a gram-negative diplococcus that colonizes genital, rectal, and oral mucosa^1^. As a strict human pathogen, *N. gonorrhoeae* is finely adapted for survival in its sole host and infects over 85 million individuals (both sexes) each year worldwide^2^. No vaccine is available, and reinfection by identical gonococcal strains is possible^1,3^. After successful colonization by gonococci (GC), the female host may experience complications arising from infection including pelvic inflammatory disease, ectopic pregnancy, infertility, and neonatal blindness caused by maternal transmission during childbirth^1^. Since the early twentieth century antibiotics have been used to treat gonococcal infections, but their effectiveness has decreased with the rise of multidrug resistance (MDR)^4^. To date, *N. gonorrhoeae* has developed resistance to sulfonamides, penicillins, tetracyclines and fluoroquinolones^1,5^. Currently, the treatment for gonococcal infections is ceftriaxone but alarmingly, ceftriaxone-resistant strains have been identified^6,7^. This consistent rise in MDR indicates the possibility of untreatable strains of *N. gonorrhoeae* and has prompted the CDC (Center for Disease Control and Prevention) to flag this bacterium as an urgent public health threat. Thus, a detailed molecular understanding of gonococcal MDR mechanisms and their regulation are crucial to inform public health decisions^6^.

*N. gonorrhoeae*, like most pathogens, contains a number of molecular mechanisms that lead to antibiotic resistance including β-lactam antibiotic resistance by antibiotic modifying proteins^8^, ribosome protective proteins^9^, and reduced permeability of cell walls by outer membrane proteins^5,10^. Significantly, the direct export of antibiotics and toxic molecules by multidrug efflux pumps is a major contributor to gonococcal survival and MDR. Multidrug efflux pumps decrease the cytosolic and periplasmic concentration of antibiotics and cytotoxins by pumping them into the extracellular milieu^11^. These systems can recognize and efflux a wide variety of chemically and structurally dissimilar molecules^11^. The best characterized multidrug efflux system of *N. gonorrhoeae* is the MtrCDE tripartite system, a member of the RND family^1^. Studies investigating MDR in *N. gonorrhoeae* report significant increase in gonococcal susceptibility to antibiotics when the MtrCDE system is impaired or deleted^12^. Overexpression of the MtrCDE efflux system is a leading cause of gonococcal resistance to hydrophobic agents (HAs), fatty acids, and even human antimicrobial peptides such as LL-37^12-14^. However, the energetic cost of synthesizing efflux pumps is significant, and their expression is regulated tightly leading to optimal cell viability^15^.

The *mtrCDE* operon is regulated by MtrR, Multiple Transferable Resistance Repressor, which also acts as a global transcriptional regulator of *N. gonorrhoeae*^12^. Beyond its regulation of the expression of the *mtrCDE* operon and MtrR directly represses *rpoH*, a sigma factor that controls the stress response in gonococci. MtrR can regulate these genes by binding a degenerate consensus sequence upstream of the coding regions thus blocking access to the promotor by RNA polymerase^12,16^. Upon binding to cytotoxic molecules and likely sensing oxidative stress, MtrR undergoes a conformational change to relieve repression of its target genes (Figure 1A)^13^.

**Figure 1:**
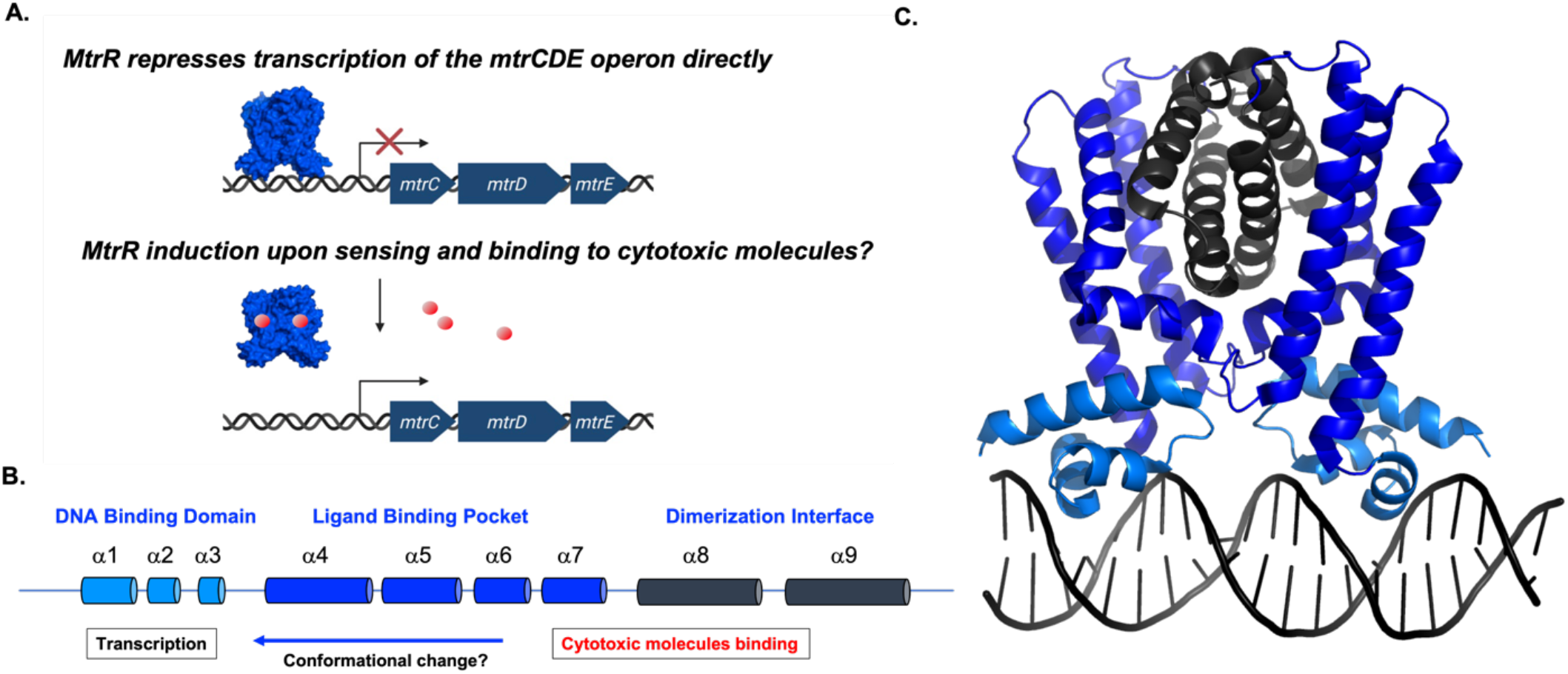
Proposed mechanism of the transcriptional regulation of the *mtrCDE* multidrug efflux pump genes by MtrR. A. MtrR bound to the *mtrCDE* operator site blocks recognition of the -35 and -10 elements by RNA polymerase thereby precluding transcription of the *mtrC, mtrD*, and *mtrE* genes, which encode for the periplasmic accessory, efflux, and outer membrane channel proteins, respectively, of the MtrCDE efflux system. The dark blue arrows depict the coding regions of each gene. The transcription start site is represented by a bent arrow. Transcriptional repression is shown as a red X, cytotoxic molecules are represented as red spheres, and MtrR is represented as a blue surface. B. The entirely helical structure of MtrR is shown as cylinders and the biochemical and biological functions of each helix is describe. C. Structure of homodimer MtrR bound to the *mtrCDE* operator site with the helices colored as in panel B (PDB: 7JU3) ^18^.

A member of the TetR family, MtrR is entirely alpha helical and comprises a helix-turn-helix (HTH) DNA binding motif and a C-terminal ligand binding/dimerization domain (Figure 1B-C)^13^. Mutations in the *mtrR* gene are present in many clinical isolates resistant to antimicrobials and antibiotics including penicillin, azithromycin, and tetracycline as well as HA’ s^17^. Recent structural and biochemical studies on the MtrR-*rpoH* and MtrR-*mtrCDE* operator complexes reveal the basis of loss of function mutations in the HTH DNA binding motif that abolish regulation of these genes by MtrR^18^.

Despite these initial studies, little is known about the induction and ligand recognition mechanisms of MtrR by innate human inducers. Previous work from our laboratory has shown that MtrR binds and is induced by bile salts chenodeoxycholate (CDCA) and taurodeoxycholate (TDCA), found at extra-urogenital GC infection sites^13^. However, physiologically relevant inducers of MtrR that are found at urogenital sites, which form the vast majority of the sites of gonococcal infection, are not known, posing a significant gap in our understanding of how MtrR contributes to MDR and gonococcal survival. To address this critical lack of knowledge, we identified several physiologically relevant steroid hormones as inducers and carried out a series of structural and biochemical experiments to elucidate their binding and induction mechanisms of MtrR.

## Results

### MtrR induction by Progesterone, Testosterone, β-Estradiol, and Ethinyl Estradiol

We first investigated the ability of MtrR to bind selected ligands. Because bacterial regulatory proteins of efflux systems are commonly induced by the substrates of the corresponding pump, allowing increased expression and resistance under cytotoxic stress, we hypothesized that some ligands of MtrCDE are MtrR inducers^19-21^. Previous work has shown that mutations of the *mtrD* gene result in significant decrease in the minimal inhibitory concentration (MICs) of progesterone (STR) in *N. gonorrhoeae*^*22*^. Other physiologically relevant candidates of MtrR inducers include the gonadal steroids β-estradiol (EST) and testosterone (TES), which, in addition to STR, have been shown to arrest *N. gonorrhoeae* growth^23^. Of these three, STR is the most effective in growth inhibition, highlighting it as potent protective agent against gonococcal infection and may explain the higher occurrence of asymptomatic gonorrheal infection in women than men^23^. Mutations in the *mtrR* gene also confer increased susceptibility to STR in plated cells as well as in female mouse models^24^.

We conducted isothermal titration calorimetry (ITC) studies on each compound to determine their affinity for MtrR. These studies reveal that MtrR binds these steroids with equilibrium dissociation constants (K_d_) of ∼ 2.8 ± 0.7 µM (STR), ∼ 1.7 ± 1 µM (EST) and 5.2 ± 3 µM (TES), whilst the chemical precursor cholesterol shows no binding (Figure 2A). The stoichiometry for EST and TES is one molecule of steroid per MtrR monomer and intriguingly one molecule of STR per MtrR dimer. In addition to these three gonadal steroids, we also tested the binding of ethinyl estradiol (NDR), which is a synthetic estrogen used in hormonal contraceptives^25^, the stress hormone, cortisol, rifampicin, and azithromycin. Of these, only ethinyl estradiol binds MtrR with a Kd of 0.94 ± 1 µM. These results show the substantial specificity of MtrR for gonadal steroid hormones. To determine if these identified host ligands result in the biochemical induction of MtrR, we measured the binding affinity of MtrR for the *rpoH* and *mtrCDE* operators in the presence and absence of 125 µM STR, EST, TES, or NDR using a DNA binding fluorescence polarization-based assay (FP). These FP experiments show up to a 12-fold and 3-fold decrease in binding affinity of MtrR for *rpoH* and *mtrCDE*, respectively, in the presence of each steroid compared to binding of MtrR to these sites in the presence of 1% methanol (MeOH) alone (Figure 2B). These data support the hypothesis that gonadal steroids are physiologically relevant inducers of MtrR.

**Figure 2:**
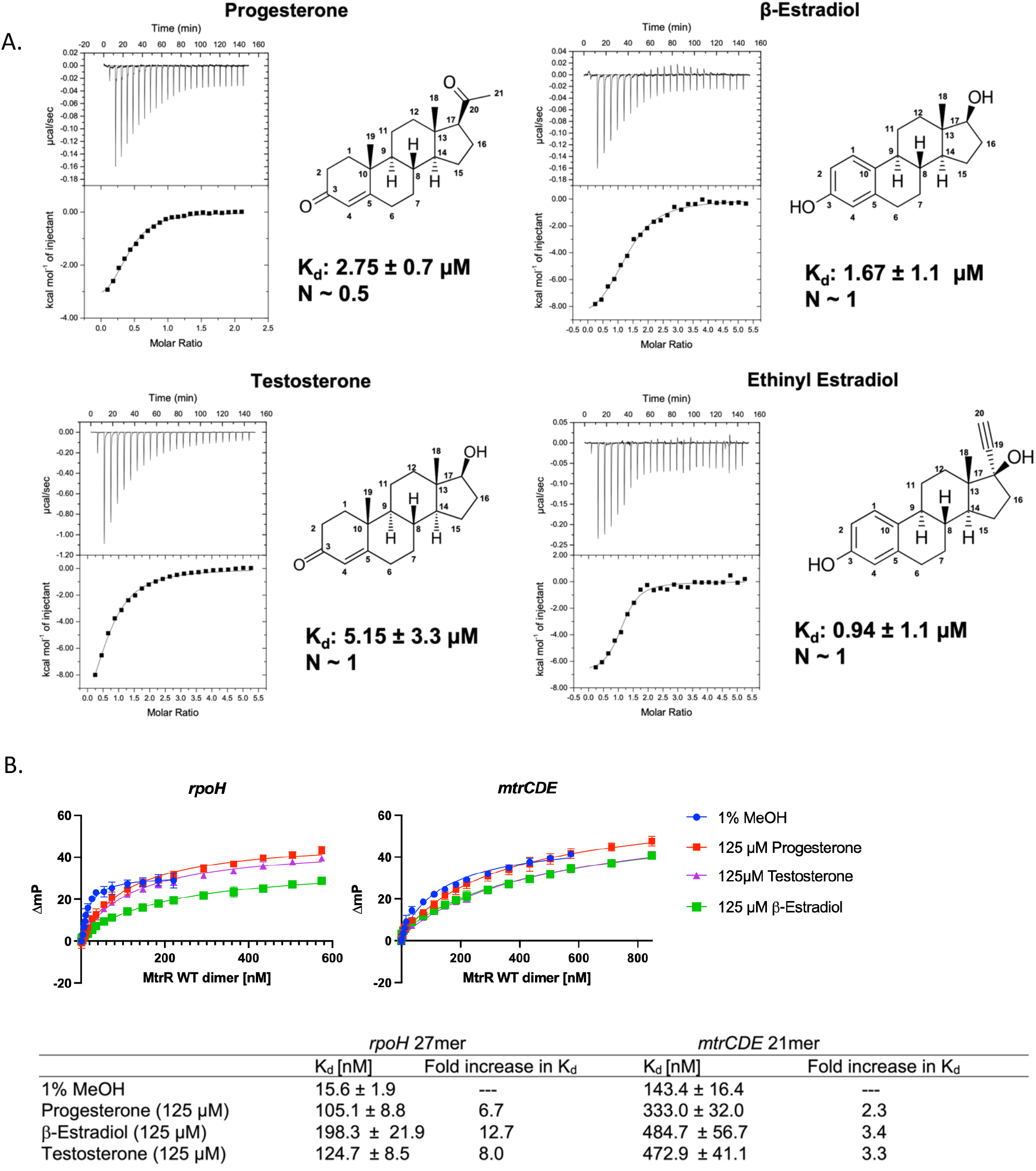
MtrR binding to the steroids Progesterone, β-Estradiol, Testosterone, and Ethinyl Estrogen. A. Isothermal titration calorimetry thermograms and binding isotherms of each steroid and ethinyl estradiol. B. Fluorescence polarization-based DNA binding isotherms reveal significant decreases in the binding affinity of MtrR for the *rpoH* and *mtrCDE* operators in the presence of each steroid.

### Structures of MtrR-Progesterone, Testosterone, β-Estradiol and Ethinyl Estrogen Complexes

We carried out crystallographic studies to determine the binding mechanisms and structural basis of induction of MtrR bound to progesterone, testosterone, β-estradiol, and ethinyl estrogen. The structures of MtrR bound to cognate DNA and an apo form have been previously solved by X-ray crystallography in our laboratory^13,18^. The structure of MtrR bound to EST was determined at a resolution of 2.37 Å using *de novo* phasing by single wavelength anomalous dispersion after selenomethionine substitution of the protein. Structures of MtrR bound to STR, TES, and NDR were subsequently solved by molecular replacement at resolutions of 2.31, 2.22 and 3.2 Å, respectively. The *R*_*work*_ and *R*_*free*_ values for each structure are 19.2% and 24.4% (MtrR+EST), 20.8% and 26.1% (MtrR+TES), 21.1% and 25.1% (MtrR+STR), and 20.1% and 27.7% (MtrR+NDR), respectively. The asymmetric unit (ASU) of all crystals contained two homodimers. The root mean square deviation (RMSD) of the pairwise alignments of all corresponding Cα of the dimers within each ASU is 0.29 Å, 0.30 Å, 0.31 Å, and 0.48 Å for the MtrR-EST, MtrR-TES, MtrR-STR and MtrR-NDR structures, respectively.

MtrR is composed of 210 residues and exists as a dimer of 48 kDa as confirmed by gel filtration chromatography.^13,18^ Consistent with the apo structure of MtrR, the first 8 to 10 residues and a stretch of 5 to 11 residues between helices α4 and α5 are not observed in the crystal structures^13^. In all structures, the last observed carboxy-terminal residue is 209 or 210. In agreement with previous structures and other TetR family members, these structures are comprised of homodimers, composed of nine α-helices. The residues of each α-helix are: α1: 10–26, α2: 33– 39, α3: 44–50, α4: 53–78, α5: 84–102, α6: 103–111, α7: 123–148, α8: 158–180, α9: 185–203 (Figure 3A). The first three helices form the DNA binding domain which contains a HTH motif, Helices α8-α9 form the majority of the dimerization interface^13,18^. The “induced” conformation of MtrR bound to each steroid is structurally identical as the RMSDs of the alignment of all Cα atoms of each dimer in all steroid-bound structures reveals that the conformations are essentially the same: MtrR+EST:MtrR+TES = 0.43 Å, MtrR+EST:MtrR+STR = 0.22 Å, MtrR+TES:MtrR+STR = 0.37 Å, MtrR+EST:MtrR+NDR = 0.46 Å, MtrR+NDR:MtrR+TES = 0.50 Å, and MtrR+NDR:MtrR+STR = 0.47 Å (Figure 3A,B).

**Figure 3:**
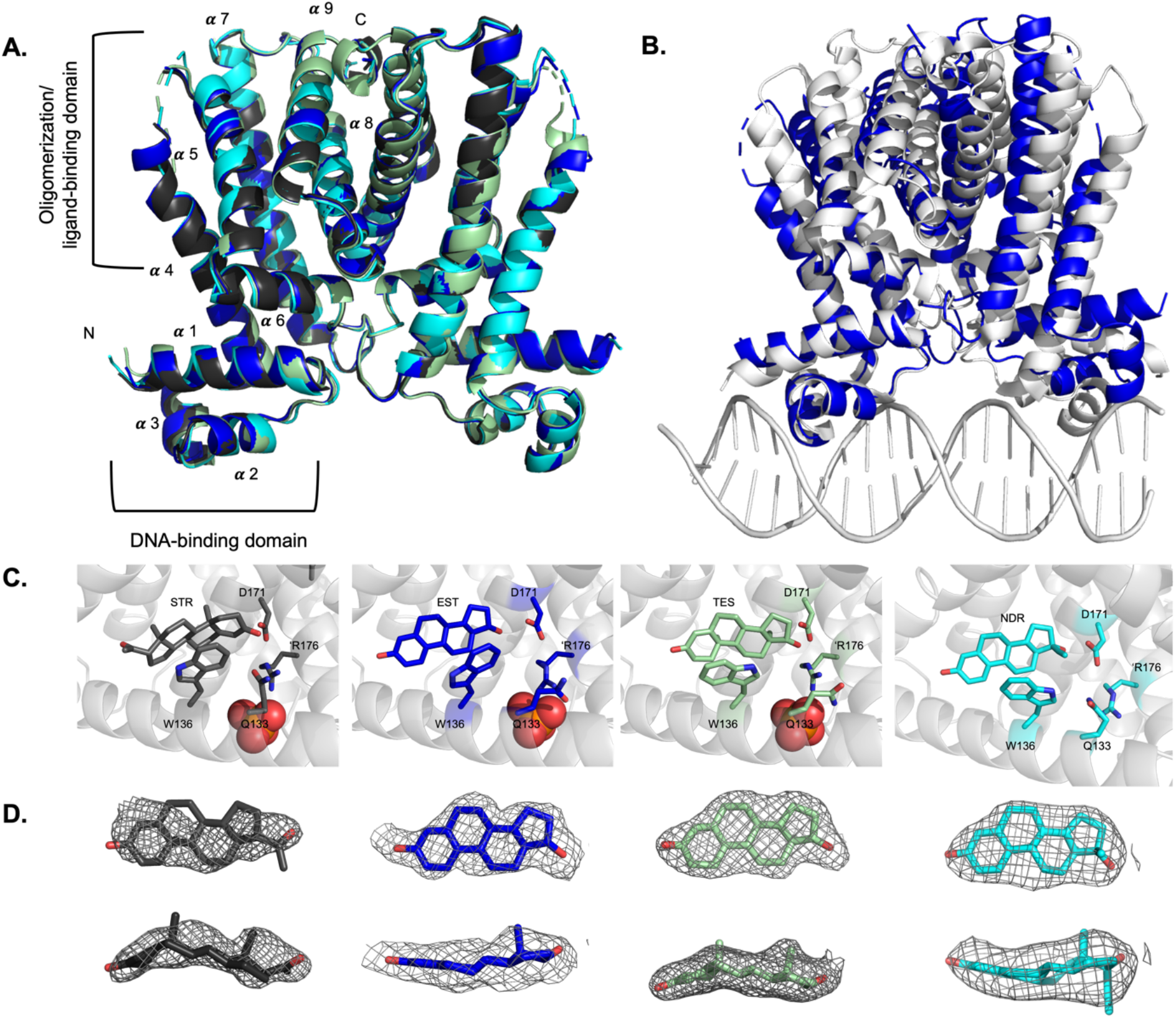
MtrR binding and induction by gonadal steroids and ethinyl estrogen. A. Overlay of hormonal steroid-bound MtrR structures with progesterone (charcoal), β-estradiol (blue), testosterone (pale green), and ethinyl estrogen (cyan). The helices of one subunit are labelled B. Overlay of MtrR bound to β-estradiol (blue) and MtrR bound to the *mtrCDE* operator site (white). C. View of MtrR binding to STR, EST, TES, and NDR. Selected binding residues are displayed as sticks. D. Electron density maps of each ligand in the MtrR ligand binding pocket looking both parallel and perpendicular to the four membered ring of each steroid. Oxygen atoms colored red. The composite Fo-Fc omit map contoured to 2.0 σ; is shown in gray mesh.

The MtrR-steroid crystal structures reveal that MtrR binds one ligand per protomer as observed in many TetR family members, including the founding member TetR, which binds one molecule of Mg^2+^-tetracycline per monomer^20^. However, not all members of the TetR family bind one molecule per protomer including QacR, which notably binds one molecule per dimer^19^. The stoichiometry of MtrR bound to EST, TES and NDR is 1 ligand:1 protomer. On the other hand, only one STR is unambiguously assigned to bind the MtrR dimer. Inspection of the binding pocket of the non STR bound-protomer reveals poor density that is possibly CAPS, a component of the crystallization buffer or a second weakly bound STR. The second dimer in the asymmetric unit contains a progesterone in each binding pocket. The stoichiometry for this MtrR-STR complex is inconsistent between the ITC experiments (n = 0.5) and the structures (n = 1). This could be in part because MtrR is crystallized in an excessive concentration of steroids which does not reflect physiological binding, the apparent progesterone concentration in the ITC experiments is not correct, or that we are observing partial occupancy of each binding site in the crystal structure.

It should be noted that the electron density for STR was the weakest, possibly indicating more than one binding pose for this steroid. β-estradiol and testosterone are positioned with the five membered ring pointing towards the dimerization surface whilst progesterone is positioned with the five membered ring positioned towards the lateral side of the protein (Figure 3 C,D). This indicates a binding pocket that is fluid allowing different ligands to bind in multiple orientations, a finding which is consistent with other TetR multidrug binding proteins such as QacR of *Staphylococcus aureus* and RamR of *Salmonella enterica* ^*19,26*^.

The ligand binding pockets of each protein-steroid complex is similar with volumes of ∼1040 Å^3^ (EST), ∼1130 Å^3^ (TES), ∼1000 Å^3^ (STR), and ∼900 Å^3^ (NDR) and consists of a tunnel between helices 4 and 7 that leads to a cavity surrounded by helices 4-9^27^. The ligand binding pocket is mainly comprised of hydrophobic residues with several amino acid residues in proximity to the steroids to provide van der Waals contacts. Notably, the aromatic side chain of W136 stacks over the rings of each steroid (Figure 3C). Interestingly, residue W136 in the MtrR-TES structure is found in two conformations but maintains van der Waals contacts to the steroid in each pose. Analysis of the composite omit maps of the MtrR-TES crystal structure reveal that these conformations are distributed equally in the asymmetric unit indicating that there is no preference for either indole ring conformation. No positively charged side chains are found in the core of the binding pocket, however three residues, K167, R176’ (‘ indicating the residue is from the other protomer of the dimer), and R137 coordinate a phosphate ion outside of the ligand binding pocket in all subunits. Residues W136 and R176 were previously shown to be important in binding the bile salt chenodeoxycholate^13^. One negatively charged amino acid residue, D171 on helix α8 near the dimerization surface is positioned ∼3.4 Å from the hydroxyl group on the five membered ring of testosterone and estradiol and ∼3.8 Å away from the carbonyl on C3 of progesterone. Due to the proximity of the side chain of D171 and the hydroxyl groups of estradiol and testosterone, this indicates an important role in MtrR ligand specificity. In addition to D171, the side chain of glutamine residue, Q133, takes multiple conformations that allow it to engage in hydrogen bonds to the carbonyl 1 of progesterone and the hydroxyl group of testosterone (Figure 3C).

Superposition of the structures of steroid-bound MtrR and MtrR bound to the *rpoH* operator sequence reveal a significant conformational change in the DNA binding domain (Figure 3B). The center-to-center distance between the “recognition” helix of the HTH motifs in the DNA bound structure is 36.8 Å. This distance is increased to 45.0, 45.2, 45.6, and 45.2 Å for MtrR bound to STR, TES, EST, and NDR, respectively. The RMSD for the overlay of these structures to the MtrR bound DNA is 1.10 (EST:DNA), 1.68 (TES:DNA), 0.96 (STR:DNA), and 1.98 (NDR:DNA). The alignment of MtrR-EST and MtrR-DNA structures reveals shared three major movements that result in the induction of the protein^28^. The first movement is a 20.8° rotation of the DNA binding domain (α1 through α3), which shifts up and away from the DNA major groove in the MtrR-EST structure (Figure 3B). This rotation results in the increased center-to-center distance of the recognition helix of the HTH domains from 36.8 Å in the DNA-bound form to ∼45 Å in the steroid-bound forms. The second significant movement is a 10° rotation and 4.5 Å shift upward of helix α4. This translation of helix α4 coincides with the upward movement of the HTH domain, away from the DNA, as well as a widening of the ligand binding pocket in the induced form. Last, the flexibility in the loop formed by residues 114-122 allows for a 17.2° outward rotation in helix α7. This movement also aids in the enlargement of the ligand binding pocket as well as a movement of residue W136 which is positioned to obstruct the entrance to the ligand binding pocket in the DNA bound form. Previously the structure determination of an “apo” structure of MtrR revealed that it took the identical induced confirmation likely due to the presence of CAPS in the ligand binding pocket^13^. Superposition of the “apo” structure and the steroid-bound structures results in an RMSD of 0.35 Å (apo-EST), 0.36 Å (apo-TES), 0.28 Å (apo-STR), and 0.54 Å (apo-NDR).

### Ligand binding to MtrR

To confirm the modes of binding of each steroid in the ligand pocket, we made single point mutations in the binding pocket of several selected residues and tested the ability of each substituted protein to bind the four steroids (Figure 4A). Residue D171 on helix α6 seemed to be the particularly important for binding and ligand specificity because its carboxy side chain was positioned within proximity to the hydroxy groups on the five-membered ring of TES, EST and NDR. The point mutation D171A was made and binding to each steroid was determined using ITC. Additionally, based on our previously results that showed residues W136 and R176 were important for bile salt binding^13^, we made and tested the binding properties of two additional mutants, W137L and R176E. We also tested the Q133A substitution because of its observed side chain flexibility that allowed the carboxamide group to interact with testosterone and progesterone. Of all mutations, D171A affected MtrR binding most significantly resulting in a 2.9-fold in K_d_for progesterone binding, a 7.0-fold decrease in testosterone affinity, and no observable binding to β-estradiol or ethinyl estradiol. Thus, this residue is important for ligand specificity, which is further buttressed by the finding that neither cholesterol nor cortisol, which do not have a polar atom in a similar position and do not bind to MtrR. The W136 mutant bound β-estradiol and testosterone similar to wild type but binding to progesterone was affected. The K_d_of MtrR W136L to progesterone is increased by 40-fold compared to WT. MtrR R176E binds testosterone and β-estradiol similarly to WT but binds progesterone with an increased K_d_of ∼3 fold. MtrR Q133A bound all ligands with K_d_values comparable to WT. Intriguingly, MtrR Q133A bound to progesterone in an endothermic reaction, unlike all other binding events, which are exothermic.

**Figure 4:**
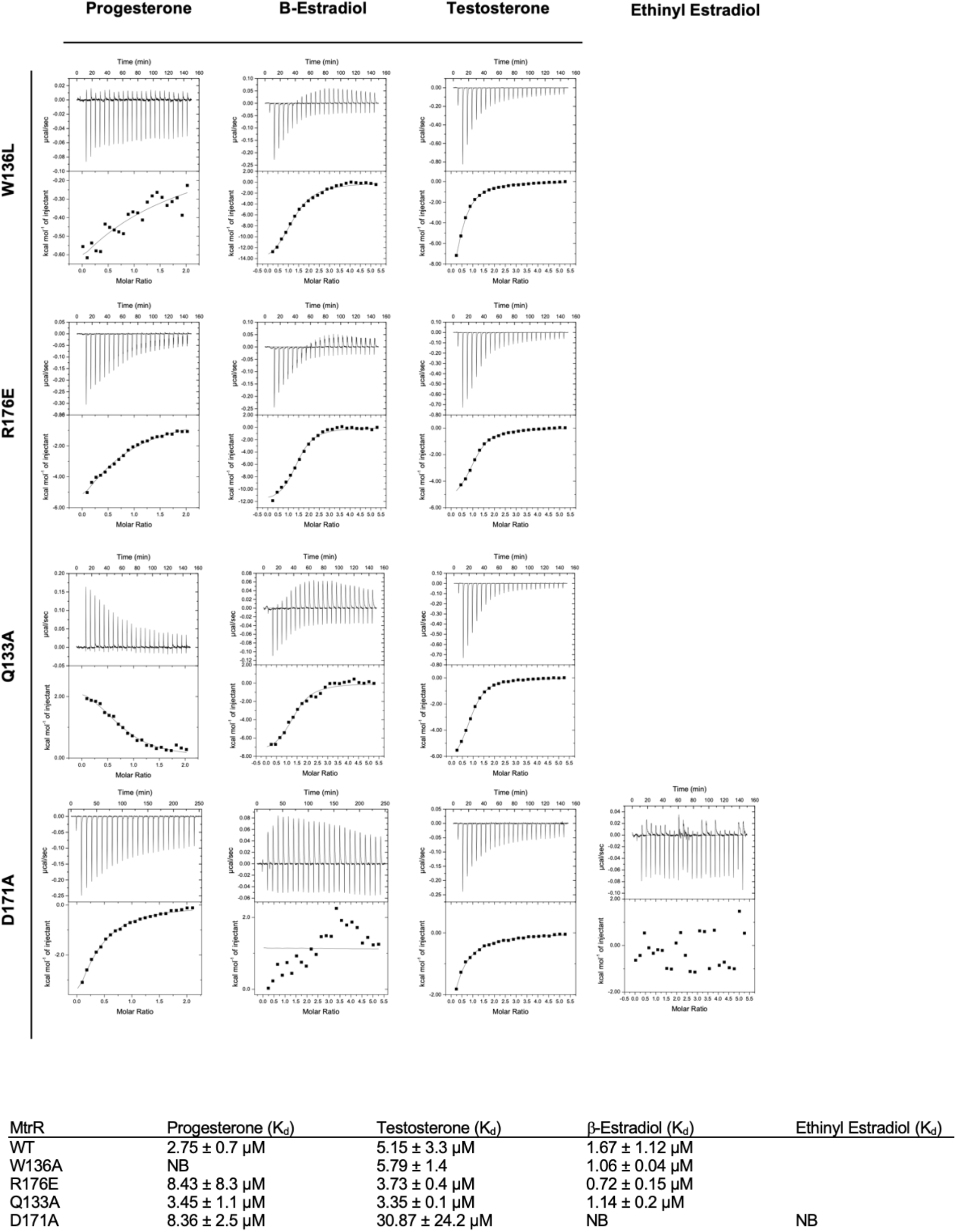
Characterization of the MtrR ligand binding pocket: Isothermal titration calorimetry thermograms and resulting binding isotherms for binding reactions between MtrR W136L, R176E, Q133A, or D171A with progesterone, β-estradiol, testosterone, and ethinyl estradiol.

## Discussion

*Neisseria gonorrhoeae* is a human pathogen and the aetiological agent of the sexually transmitted infection, gonorrhea. Multiple strains of *N. gonorrhoeae* are multidrug-resistant (MDR), and this bacterium has been identified as an urgent public health threat. Understanding the mechanism(s) of MDR in pathogenic bacteria such as *N. gonorrhoeae* is paramount to combating the prospect of untreatable infections. The overexpression of multidrug efflux pumps that can export myriad antibiotics or other cytotoxic molecules from the cell is an important mechanism for *N. gonorrhoeae* survival and the development of MDR. By tightly regulating the expression of these efflux pumps, the bacteria can respond quickly to drug treatment or attacks by the host whilst minimizing the energetic cost of efflux pump biosynthesis. Here, we identify the gonadal steroid hormones testosterone, β-estradiol, and progesterone, as well as ethinyl estrogen, a component of hormonal birth control pills, as physiologically relevant inducers of the MtrR, the MtrCDE efflux and RpoH stress response system. During gonococcal colonization of the urogenital tract, this bacterium will encounter potentially toxic local concentrations of these steroidal hormones but induces the MtrCDE system allowing *N. gonorrhoeae* to survive in the presence of these compounds by their efflux.

Additionally, we report the structures of MtrR bound to these steroids to reveal the ligand-recognition and induction mechanisms and structural consequences of ligand binding. These structures reveal a highly hydrophobic/aromatic binding pocket of MtrR and the presence of a key aspartate residue that accommodates multiligand binding. Site directed mutagenesis of selected residues within the ligand binding pocket reveal that D171 is perhaps the most critical contributor to hormonal steroid binding. Additionally, the interaction between D171 and a polar atom of each steroid appears to be a key determinant in discriminating against other steroids such as cholesterol and cortisol, which are lacking an appropriate proximal polar group. Such a binding pocket is consistent with other multidrug binding proteins such as QacR from *Staphylococcus aureus*^*19*^, TtgR from *Pseudomonas putida*^*29*^, CmeR from *Campylobacter jejuni*^*30*^and AcrR from *Eschericia coli*^*31*^, the drug-binding sites of which consist of mostly hydrophobic and aliphatic residues with few polar residues. The structures of the MtrR-steroids described here, represent the induced form of MtrR as the center-to-center distance between the two helix-turn-helix DNA binding motifs is not suitable for recognition of the nucleobases of two consecutive major grooves when compared to the DNA bound conformation of MtrR^18^. Such an increase in this distance is consistent with other induction mechanisms of TetR family members^19,32-35^. Although members of the TetR family show strong structural homology in their DNA binding domains, the ligand binding pockets vary in composition and volume. For example, the ligand binding pocket of QacR, is a bit unusual for an multiligand binding protein as it utilizes glutamates that become available upon ligand binding due to a conformational change, as well as polar residues to facilitate the binding of multiple drugs, but maintains critical aromatic interactions with each compound^19^. Binding of bile salts to RamR is facilitated by four hydrogen bonding residues, whilst relying only on π-π stacking to bind compounds such as dequalinium, ethidium bromide, and rhodamine 6G^26,36^. Here, we show that MtrR utilizes only one polar residue, D171, to bind specifically to gonadal steroids.

In conclusion, we have elucidated the inducer binding mechanism of MtrR, a transcriptional regulator that is a key player in *Neisseria gonorrhoeae’ s* multidrug resistance arsenal, by hormonal steroids. Significantly, the induction of the *mtrCDE* efflux genes and the gene encoding the *rpoH* stress response sigma factor by ethinyl estrogen indicates that steroid-based hormonal contraceptives might be utilized by *Neisseria gonorrhoeae* to help colonize its human host.

## Materials and Methods

### Protein overexpression and purification

The gene encoding *mtrR* is codon optimized (GenScript) and synthesized for expression in *E. coli*. The gene was subsequently subcloned into the ampicillin resistant pMCSG7 vector in-frame with a T7 promotor, N-terminal Hexa-histidine tag, and tobacco etch virus (TEV) protease cleavage site. The plasmid was transformed into Rosetta Gami B(DE3) pLysS cells. The cells were grown in Luria-Bertani broth containing ampicillin, chloramphenicol, and tetracycline at 37°C. Cultures were grown to an optical density (OD_600_) of 0.5-0.6 absorbance units and expression of MtrR was induced by the addition of 1 mM isopropyl-β-D-thiogalactopyranoside (IPTG) and grown for an additional 3 h before harvesting. Cells were centrifuged, reconstituted in buffer containing 20 mM Tris-HCl pH 8.0, 200 mM NaCl, 10% glycerol, and 1 mM tris-2-carboxyethyl phosphine hydrochloride (TCEP) and lysed by sonication. MtrR was purified from clarified lysate by Ni^2+^-nitrilotriacetic acid (Ni-NTA) affinity chromatography followed by cleavage of the hexa-histidine tag by TEV digestion. MtrR was further purified by size exclusion chromatography (S200 column). Selenomethionine modification of MtrR was achieved using the methionine-inhibitory pathway and purified as above^37^.

### Crystallization and data collection

Crystals were grown using the hanging drop vapor diffusion method at 25 °C. MtrR was co-crystalized with 2.5 mM progesterone, β-estradiol, testosterone, or ethinyl estradiol with 5% MeOH. Crystals formed within two weeks in crystallization solution containing 1200 mM sodium phosphate monobasic, 800 mM potassium phosphate dibasic, 100 mM CAPS/ NaOH pH 10.5, and 200 mM lithium sulfate. Crystals were cryoprotected in this buffer and 25% glycerol for flash freezing in liquid nitrogen. Data were collected at the Advance Light Source (ALS) on beamline 5.0.2 (MtrR-EST and MtrR-TES) 5.0.1 (for MtrR-STR), or 5.0.3 (for MtrR-NDR). The wavelength for data collection was 0.9763 (MtrR-EST), 0.9786 (MtrR-TES), 0.9774 (MtrR-STR), and 0.9765 (MtrR-NDR). Data were collected at 100 K. The structure of the MtrR-β-estradiol complex was determined using SAD methods via selenomethione-substituted MtrR for initial phasing. This structure was used for molecular replacement for MtrR bound to progesterone, testosterone, and ethinyl estrogen. Iterative cycles of model building were done in COOT^38^ and refinement using Phenix refine^39-41^. The stereochemistry was excellent with the percent Ramachandran favored for each structure is 98.82 (MtrR-EST), 98.02 (MtrR-TES), 98.25 (MtrR-STR), and 87.86 (MtrR-NDR.) There were no Ramachandran outliers in any structure except for MtrR bound to NDR which had the lowest resolution. Clash scores for all structures were less than or equal to 6 except MtrR bound to NDR, which had a clashscore of 10.83. Mean I/sigma(I) values for the highest resolution shell were 0.7 (MtrR-EST), 2.2 (MtrR-TES), 2.3 (for both MtrR-STR and MtrR-NDR). Selected data collection and refinement statistics are listed in Table 1.

**Table 1:**
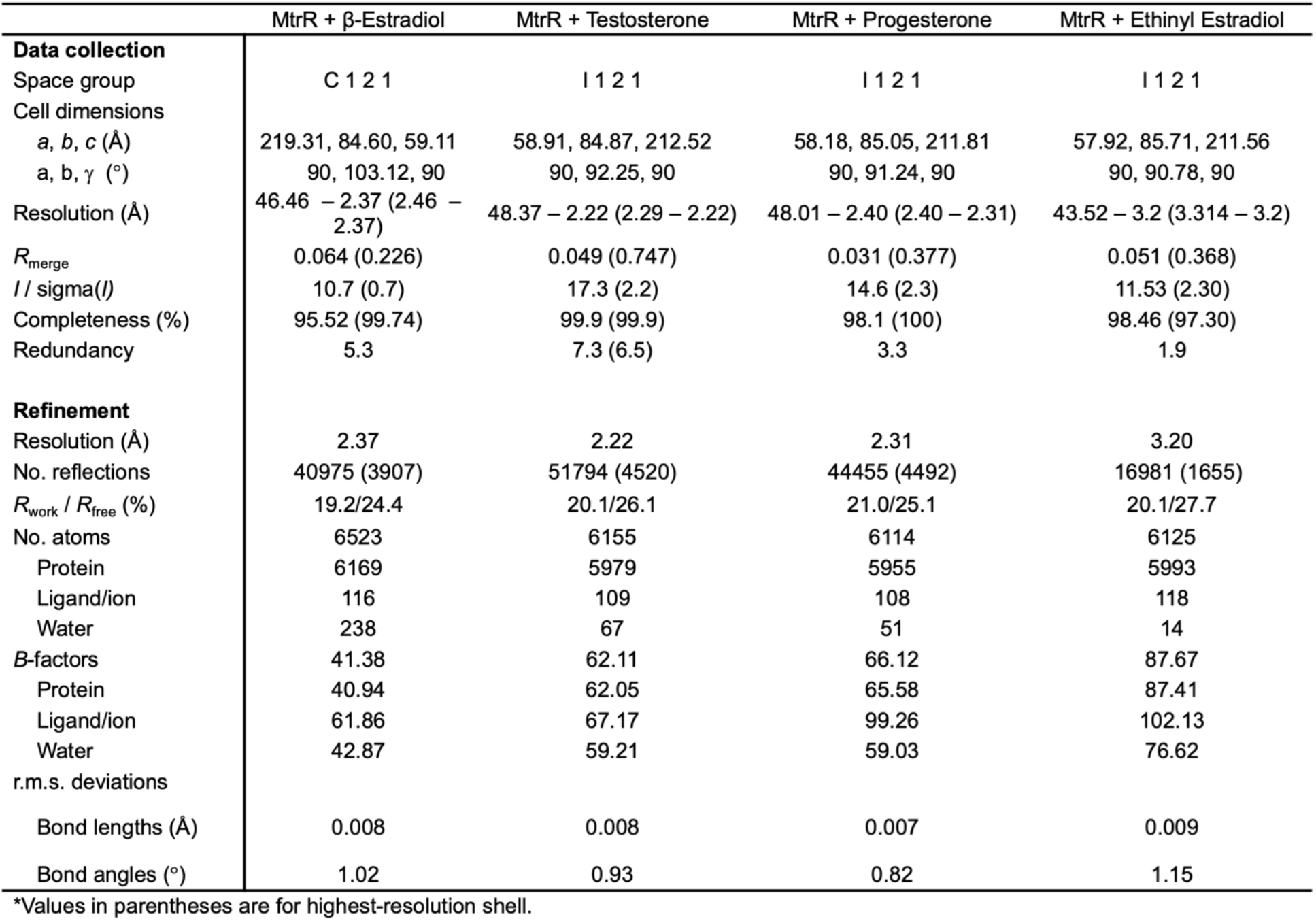
Data collection and refinement statistics.

### Isothermal Titration Calorimetry

Purified MtrR was concentrated to 20 µM in purification buffer plus 1% MeOH. The addition of 1% methanol was necessary to dissolve each steroid in the purification buffer. Titrations with 20 or 5 µM MtrR in the sample cell and 125, 200, or 500 µM compound in the syringe were performed using a VP-ITC microcalorimeter (Microcal Inc.). ITC experiments were conducted at 25 °C with a stirring speed of 199 rpm. Germane blanks were subtracted, and the data were analyzed using ORIGIN 7.0.

### Fluorescence polarization-based DNA-binding assays

Fluorescence polarization DNA-binding data were collected with a Panvera Beacon 2000 fluorescence polarization system (Invitrogen) and analyzed with Prism (Graphpad Software). Purified MtrR was buffer exchanged into “DNA” binding buffer (100 mM NaCl, 20 mM Tris-HCl pH 7.5, 2.5% glycerol, 1 mM TCEP). MtrR was titrated into a binding buffer solution that contained 1 nM 5^’^ -fluoresceine-labelled DNA (containing either the *rpoH* or *mtrCDE* operator site), 1 µg of poly(dI-dC) DNA, 1 µg bovine serum albumin (BSA) and 125 µM progesterone, testosterone, estradiol or ethinyl estradiol resulting in the final methanol concentration of 1%. BSA and dI-dC are included as carriers and to control for nonspecific DNA binding. Samples were excited at 490 nm and polarized emissions measured at 530 nm. Controls of DNA ± methanol, ligands, and MtrR were run separately. The binding affinity of protein for DNA was determined using the following equation: P = P_f_+ (P_b_ – P_f_) [M] / (K_d_ + [M]) where P is the measured polarization, P_f_ is the polarization of free DNA, P_b_ is the polarization of the completely bound DNA, K_d_ is the equilibrium dissociation constant, and M is the concentration of protein. P_b_ and K_d_ were determined by nonlinear least-squares analysis. At least three independent experimental measurements were averaged for each reported binding constant.

### Accession numbers

The coordinates and structure factors for the MtrR-STR, MtrR-EST, MtrR-TES, and MtrR-NDR complexes have been deposited in the RCSB Protein Data Bank with the accession codes 8FW8, 8FW0, 8FW3, and 8SSH respectively.

## Acknowledgements

The authors would like to thank the beamline scientists at ALS BL 8.3.1 and BL 5.0.2 for their help with data collection. In addition, we acknowledge the Advanced Light Source at Lawrence Berkeley National Laboratory. National Institutes of Health [R35 GM130290 to M.A.S., R05 AI048593 to R.G.B., R37 AI021150-35 to W.M.S.]; W.M.S. is the recipient of a Senior Research Career Scientist Award from the Biomedical Laboratory Research and Development Service of the U.S. Department of Veterans Affairs; The Advanced Light Source is supported by the Director, Office of Science, Office of Basic Energy Sciences, Material Sciences Division, of the U.S. Department of Energy [DE-AC03-76SF00098]. Funding for open access charge: Duke University School of Medicine.

## Author Contributions

G.M.H. and R.G.B. designed the experiments and analyzed the biochemical data. G.M.H., R.G.B. and M.A.S. analyzed the structural data. G.M.H. and G.A.B. generated purification constructs. G.M.H. purified proteins, determined their structures and performed biochemical characterizations. M.A.S. and G.M.H. collected x-ray crystallography data and M.A.S. provided x-ray crystallography consulting and experimental input. G.M.H. and R.G.B wrote the manuscript. We thank J.C.A. and W.M.S for providing comments on the manuscript and insights regarding the potential biological implications of our work. We also thank J.R.P. for insights into steroid and pathogen biology and for providing testosterone for the project. All authors have read and approved the manuscript.

